# Bridging verbal coordination and neural dynamics

**DOI:** 10.1101/2024.04.23.590817

**Authors:** Isaïh Schwab-Mohamed, Manuel R. Mercier, Agnès Trébuchon, Benjamin Morillon, Leonardo Lancia, Daniele Schön

**Affiliations:** Aix Marseille University, Inserm, INS, Inst Neurosci Syst, Marseille, France; Aix-Marseille Univ, Institute of Language, Communication and the Brain, France; APHM, Hôpital de la Timone, Service de Neurophysiologie Clinique, Marseille, France; Aix Marseille University, CNRS, Laboratoire Parole et Langage (LPL), Aix-en-Provence, France

## Abstract

Our use of language, which is profoundly social in nature, essentially takes place in interactive contexts and is shaped by precise coordination dynamics that interlocutors must observe. Thus, language interaction is highly demanding on fast adjustment of speech production. Here, we developed a real-time coupled-oscillators virtual partner that allows - by changing the coupling strength parameters - to modulate the ability to synchronise speech with a virtual speaker. Then, we recorded the intracranial brain activity of 16 patients with drug-resistant epilepsy while they performed a verbal coordination task with the virtual partner (VP). More precisely, patients had to repeat short sentences synchronously with the VP. This synchronous speech task is efficient to highlight both the dorsal and ventral language pathways. Importantly, combining time-resolved verbal coordination and neural activity shows more spatially differentiated patterns and different types of neural sensitivity along the dorsal pathway. More precisely, high-frequency activity in left secondary auditory regions is highly sensitive to verbal coordinative dynamics, while primary regions are not. Finally, while bilateral engagement was observed in the high-frequency activity of the IFG BA44—which seems to index online coordinative adjustments that are continuously required to compensate deviation from synchronisation— interpretation of right hemisphere involvement should be approached cautiously due to relatively sparse electrode coverage. These findings illustrate the possibility and value of using a fully dynamic, adaptive and interactive language task to gather deeper understanding of the subtending neural dynamics involved in speech perception, production as well as their interaction.

## Introduction

Language has been most frequently studied by separately assessing perception and production in an isolated context, in contrast to the interactive context that characterizes best its daily usage. The use of language in an interactive context, particularly during conversations, calls upon numerous predictive and adaptive processes (Pickering & Gambi, 2018). Importantly, analyses of conversational data have highlighted the phenomenon of interactive alignment (Pickering & Garrod, 2004), illustrating that during a verbal exchange interlocutors tend to imitate each other and to align their linguistic representations on several levels including phonetic, syntactic, or semantic (Garrod & Pickering, 2009). This is an unconscious and dynamic phenomenon that possibly renders exchanges between speakers more fluid (Marsh et al., 2009). It consists in mutual anticipation (prediction) and coordination of speech production, leading, for instance, to a reduction of turn-taking durations (Levinson, 2016 ; Corps et al., 2018).

Recently, in an effort to assess speech and language in more ecological contexts, researchers in neuroscience have used interactive paradigms to study some of these coordinative phenomena. These studies involved turn-taking behaviours such as alternating naming tasks (Mukherjee et al., 2019), questions and answers investigating motor preparations (Bögels et al., 2015), or manipulating the turn predictability in end-of-turn detection tasks (Magyari et al., 2014; for a review see, Bögels et al., 2017). However, the synchronous speech paradigm has been overlooked. This paradigm requires the simultaneous and synchronized production of the same word, sentence, or text between two people. Interestingly, this task is remarkably well performed without any particular training, both between a speaker and a recording as well as between two speakers (Cummins, 2002; Cummins, 2003; Assaneo et al., 2019). As in most joint tasks, individuals must mutually adjust their behaviour (here speech production) to optimize coordination (Cummins, 2009). Furthermore, synchronous speech favors the emergence of alignment phenomena, such as the fundamental frequency or the syllable onset (Assaneo et al., 2019; Bradshaw & McGettigan, 2021 ; Bradshaw et al., 2023 ; Bradshaw et al., 2024). Overall synchronous speech represents a strong interactive framework allowing a good level of experimental control. It offers several possibilities for neurophysiological investigation of both speech perception and production and is an interesting case to consider for models of speech motor control.

Synchronous speech resembles to a certain extent delayed/altered auditory feedback tasks, which involve real-time perturbations in the speech production signal (such as changes in fundamental frequency and delay). These tasks can induce speech errors as well as modulations in speech and voice features (Stuart et al., 2002; Yamamoto & Kawabata, 2014; Karlin et al., 2021). Additionally, these tasks provide insights into predictive models of speech motor control, where the brain generates an internal estimate of production and corrects errors when auditory feedback deviates from the estimate (Hickok et al., 2011; Houde & Nagarajan, 2011; Tourville & Guenther, 2011; Ozker et al., 2022; Floegel et al., 2023). Previous studies have revealed increased responses in the superior temporal regions compared to normal feedback conditions (Hirano et al., 1997; Hashimoto & Sakai, 2003; Takaso et al., 2010; Ozker et al., 2022; Floegel et al., 2020 ; see Meekings & Scott, 2021 for a review of error-monitoring and feedback control in the STG during speech production). However, synchronous speech paradigms allow for the investigation of the neural bases of coordinative behaviour rather than of error correction.

So far, the precise spectro-temporal dynamics and spatial distribution of the cortical networks underlying speech coordination remain unknown. To address this issue, we first developed a real-time coupled-oscillator virtual partner that allows - by changing the coupling strength parameters - to modulate the ability to synchronise speech with a speaker. The virtual partner (Lancia et al.,2017) and the synchronous speech task were first tested on a control group to ensure the ability of the virtual agent to coordinate its speech production in real time with the participants. In certain conditions, the agent was programmed to actively cooperate with the participants by synchronizing its syllables with theirs. In other conditions, the agent was programmed to deviate from synchronization by producing its syllables between those of the participants. As a result, participants were constantly required to adapt their verbal productions in order to maintain synchronization. Appropriate tuning of the coupling parameters of the virtual agent enabled us to create a variable context of coordination yielding a broad distribution of phase delays between the speaker and agent productions. Subsequently, we leveraged the excellent spatial sensitivity and temporal resolution of stereotactic depth electrodes recordings and acquired neural activity from 16 patients with drug-resistant epilepsy while they performed the adaptive synchronous speech task with the virtual partner.

## Materials and methods

### Control – participants

30 participants (17 women, mean age 24.7 y, range 19-42 y) took part in the study. All were French native speakers with normal hearing and no neurological disorders. Participants provided written informed consent prior to the experimental session and the experimental protocol was approved by the Institutional Review board of the French Institute of Health (IRB00003888). 5 participants (2 women) were excluded from analysis for poor signal-to-noise ratio in speech recordings.

### Patients – participants

16 patients (7 women, mean age 29.8 y, range 17 - 50 y) with pharmacoresistant epilepsy took part in the study. They were included if their implantation map covered at least partially the Heschl’s gyrus and had sufficiently intact diction to support relatively sustained language production. All patients were French native speakers. Neuropsychological assessments carried out before stereotactic EEG (sEEG) recordings indicated that they had intact language functions and met the criteria for normal hearing. In none of them the auditory areas were part of their epileptogenic zone as identified by experienced epileptologists. Recordings took place at the Hôpital de La Timone (Marseille, France). Patients provided written informed consent prior to the experimental session and the experimental protocol was approved by the Institutional Review board of the French Institute of Health (IRB00003888).

### Data acquisition

The speech signal was recorded using a microphone (RODE NT1) adjusted on a stand so that it was positioned in front of the participant’s mouth. Etymotic insert earphones (Etymotic Research E-A-R-TONE gold) fitted with 10mm foam eartips were used for sound presentation. The parameters and sound adjustment were set using an external low-latency (∼5ms) sound card (RME Babyface Pro Fs, Kim et al., 2020), allowing a tailored and temporally precise configuration for each participant. A calibration was made to find a comfortable volume and an optimal balance for both the sound of the participant’s own voice, which was fed back through the headphones, and the sound of the stimuli. The aim of this procedure was that the patient would subjectively perceive their voice and the VP-voice in equal measure. VP voice was delivered at approximately 70dB.

The sEEG signal was recorded using depth electrodes shafts with a 0.8 mm diameter containing 5 to 18 electrode contacts (Dixi Medical or Alcis, Besançon, France). The contacts were 2 mm long and were spaced from each other by 1.5 mm. The placement of the electrode implantations was determined solely on clinical grounds. Sixteen patients with a total of 236 electrodes (145 in the left hemisphere) and 2395 contacts (1459 in the left hemisphere, see Figure 1). While this gives a rather sparse coverage of the right hemisphere, we decided, due to the rarity of this type of data, to report results for both hemispheres, with figures for the left hemisphere in the main text and figures for the right hemisphere in the supplementary section. All Patients were recorded in a sound-proof Faraday cage using a 256-channels amplifier (Brain Products), sampled at 1kHz and high-pass filtered at 0.16 Hz.

**Figure 1.**
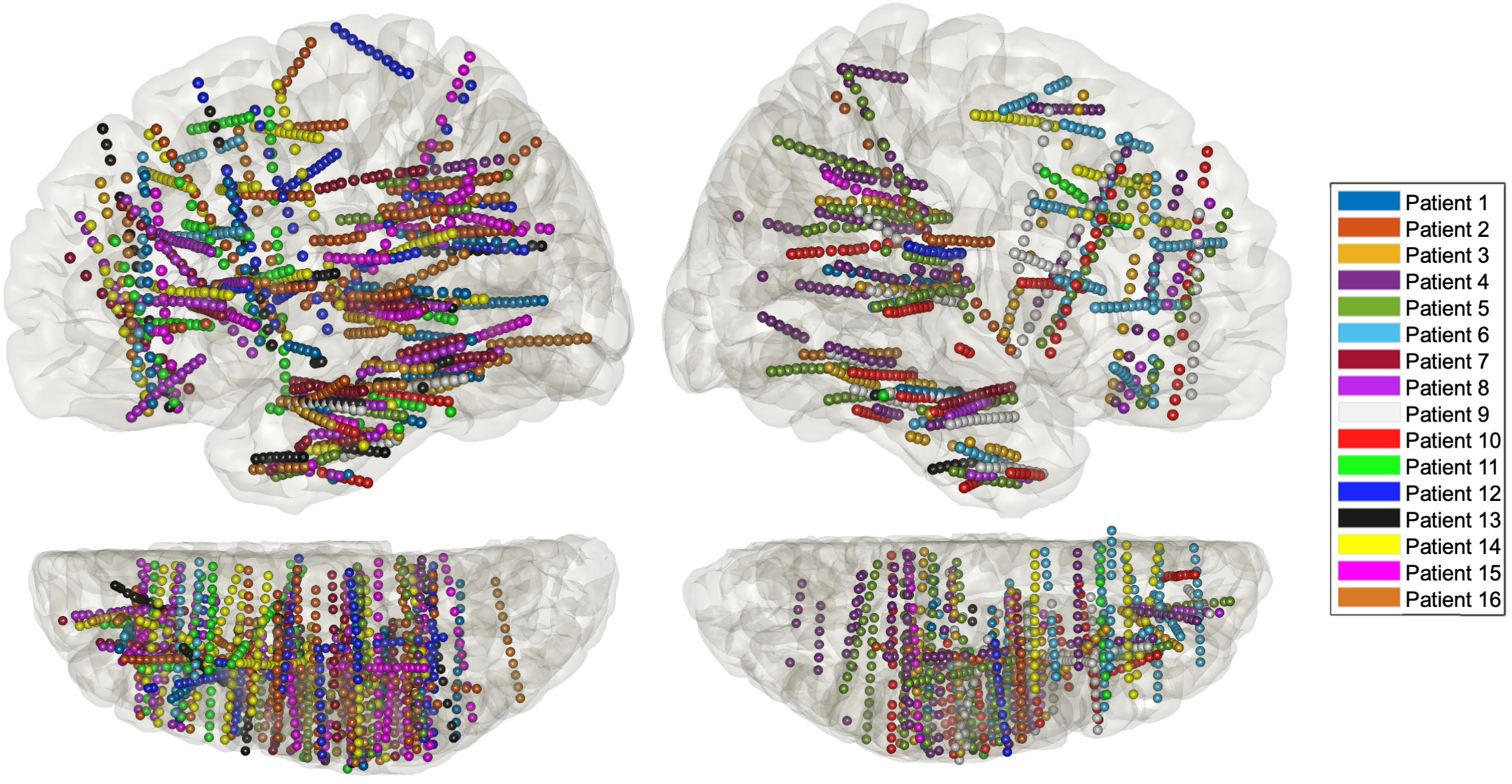
Anatomical localization of the sEEG electrodes for each patient projected in MNI space on the lateral 3D view (top) and on the top view (N=16).

### Stimuli

Four stimuli corresponding to four short sentences were pre-recorded by both a female and a male speaker. This allowed to adapt to the natural gender differences in fundamental frequency (i.e. so that the VP gender matched that of the patients). All stimuli were normalised in amplitude.

Stimuli consisted of four sentences: “papi mamie” (/*papi mami*/, grandpa grandma) / “papi m’a dit” (/*papi ma di*/, grandpa told me) / “mamie lavait ma main” (/*mami lavɛ ma mɛ”*/, grandma washed my hand) / “mamie manie ma main” (/*mami mani ma mɛ”*/, grandma handles my hand). The four sentences purposely differed in terms of number of syllables (4-4-6-6). Moreover, two of them contained deviations from an otherwise repeating phonological pattern: both in /*papi ma di*/ and in /*mami mani ma mɛ”*/, the repeated opening/closing of the lips is substituted by the formation and release of a tongue constriction. These manipulations made the sentences more or less easy to articulate (easy-medium-easy-medium). Sentence durations were 2.07, 2.11, 3.16 and 2.87s, respectively with a syllable rate of ∼3Hz.

### Experimental design

Participants, comfortably seated in a medical chair, were instructed that they would perform a real-time interactive synchronous speech task with an artificial agent (Virtual Partner, henceforth VP, see next section) that can modulate and adapt to the participant’s speech in real time.

The experiment required 3 steps (Figure 2A). First, the sentences were presented in written form and the experimenter verified that each participant could pronounce them correctly. Second, a training phase took place. This required repeating each stimulus over and over together with the VP for ∼14 seconds. More precisely, the sentence was first presented on a screen. When the participant pressed the “space” bar, the visual stimulus went off, and the VP started to « speak ». The participant was instructed to repeat the stimuli as synchronously as possible with the VP for the whole trial duration. In the training phase the VP did not adapt to the participant.

**Figure 2.**
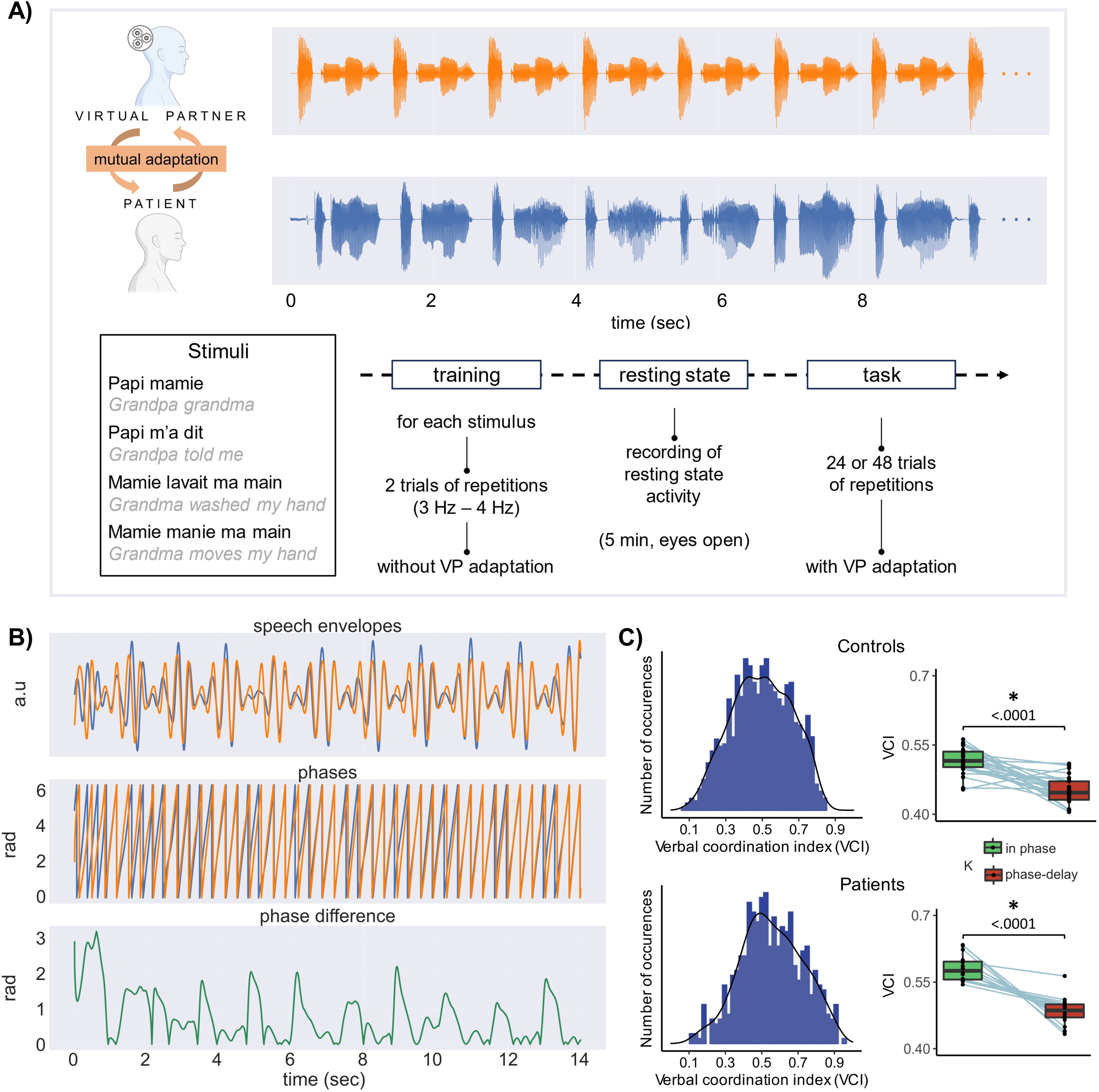
Paradigm and coordination indexes. **(A) Top:** illustration of one trial of the interactive synchronous speech repetition task (orange: virtual partner speech; blue: participant speech; stimulus papi m’a dit repeated 10 times ; only the 10 first seconds are represented). **Bottom:** the four speech utterances used in the task and the experimental procedure. **(B)** Speech signals processing stages. The top panel corresponds to the speech envelope, the second to the phase of speech envelope and the third panel to the phase difference between VP and participant speech envelopes, illustrating the coordination dynamics along one trial. **(C)** Left: distributions of verbal coordination index (phase locking values between VP and participant speech envelopes, for each trial) for all participants (top) and patients. Right: boxplots for control participants (top) and patients showing the trial-averaged verbal coordination index as a function of the virtual partner parameters (in-phase coupling vs coupling with a 180° shift).

The training allowed participants to familiarise with the synchronous speech task but also to build a personalised virtual partner model incorporating the articulatory variability of each participant (see section below).

The training was followed by a resting phase, allowing the recording of resting state activity for a period of 5 min. This time also allowed to build, for each participant, the virtual partner model.

The third step was the actual experiment. This was identical to the training but consisted of 24 trials (14s long, speech rate ∼3Hz, yielding ∼1000 syllables). Importantly, the VP varied its coupling behaviour to the participant. More precisely, for a third of the trials the VP had a neutral behaviour (close to zero coupling: k = +/- 0.01). For a third it had a moderate coupling, meaning that the VP synchronised more to the participant speech (k = - 0.09). And for the last third of the trials the VP had a moderate coupling but with a phase shift of pi/2, meaning that it moderately aimed to speak in between the participant syllables (k = + 0.09). The coupling values were empirically determined on the basis of a pilot experiment in order to induce more or less synchronization but keeping the phase-shifted coupling at a rather implicit level. In other terms, while participants knew that the VP would adapt, they did not necessarily know in which direction the coupling went. Depending on patient fatigue, a second experimental session with 24 extra trials, was proposed (for 6 of the 16 patients). The control group participant run a single session of 48 trials.

### Virtual partner (principles)

The virtual partner (henceforth VP) used for the experiment allows to generate speech (words or short utterances) while adapting it in real-time to the concurrent speech input. The VP environment is built upon the Psychtoolbox-3 program and operates within MATLAB, leveraging custom C subroutines to boost its performance. Its operation revolves around a loop whose iterations are executed at consistent intervals of Δt = 4ms. During each iteration, the program analyzes the latest segment (25ms) of speech produced by the participant and streams a portion of speech to the output device. More precisely, at each iteration of the main loop, the functioning of the VP can be described in four steps:

1. A feature vector is extracted from the last chunk of the input signal.
2. The phase value of the input signal chunk is calculated by mapping the input feature vector onto the corresponding vector of the stimulus signal and retrieving the associated phase value. This step is performed using a dynamic-time-warping algorithm (Dixon, 2005). To enhance precision, the input chunk is mapped onto several model utterances (all time-aligned with the signal used as a stimulus) that are tailored to the characteristics of the participant’s speech in the training phase.
3. A chunk of stimulus signal is chosen to be sent to the output device. The selection is guided by applying the Kuramoto (1975) equation to the difference between the phase values representing the current positions of the participant and the VP in their syllabic cycles. This enables real-time lengthening or shortening of speech chunks as needed to adjust towards the preferred phase (0 or π/2). Significantly, in applying this equation, we assign a specific value to the coupling strength parameter « k », linking the behaviour of the VP to that of the participant. Values can vary from trial to trial and could be close to 0 (k = +/- 0.01) resulting in a neutral behaviour, negative values (k = - 0.09) equivalent to a moderate coupling behaviour (tendency to synchronize), or positive values (k = + 0.09) corresponding to a moderate coupling behaviour with a phase shift of π/2 (tendency to speak in between participant syllables).
4. The selected chunk of stimulus signal is integrated into the output stream via WSOLA synthesis (Waveform Similarity Overlap-Add).

Of note, the coupling strength were chosen to be rather weak and thus do not allow to reach 0 or π/2 phase synchrony, but rather yield the desired large panel of phase delays in the VP-participant coordinative behaviour (see Figure1).

### Data analysis speech signal

Speech signals of the participants and the VP were processed using Praat and Python scripts.

First, raw speech signals were downsampled from 48KHz to 16KHz. Then speech envelope was extracted using a pass-band filter between 2.25 and 5Hz. The phase was computed using the Hilbert transform.

To quantify the degree of coordination of the verbal interaction (verbal coordination index, VCI) we computed the phase locking value using mne_connectivity python function spectral_connectivity_time with the method ‘plv’ based on the phase locking value index proposed by Lachaux et al. (1999). Phase locking was computed for each trial on speech temporal envelope (Fig 1.B), resulting in 24 or 48 verbal coordination indexes (VCI) per patient depending on the number of sessions performed. To assess the effect of the coupling parameter we computed a linear mixed-model contrasting *in-phase coupling trials and trials with a coupling set towards a 180° shift* (lmer(VCI ∼ K + (1|participant)). As expected, because of the varying adaptive behaviour of the VP, VCI varies across trials, indexing more or less efficient coordinative behaviour (Figure 1C). Moreover, in order to estimate whether the level of performance was greater than chance level, we computed, for each patient and each trial, a null distribution obtained by randomly shifting the phase between the VP and the patient speech (500 times, see Figure S1).

### sEEG signal

#### General preprocessing related to electrodes localisation

To increase spatial sensitivity and reduce passive volume conduction from neighbouring regions (Mercier et al., 2017), the signal was offline re-referenced using bipolar montage. That is, for a pair of adjacent electrode contacts, the referencing led to a virtual channel located at the midpoint locations of the original contacts. To precisely localize the channels, a procedure similar to the one used in the iELVis toolbox was applied (Groppe et al., 2017). First, we manually identified the location of each channel centroid on the post-implant CT scan using the Gardel software (Medina Villalon et al., 2018). Second, we performed volumetric segmentation and cortical reconstruction on the pre-implant MRI with the Freesurfer image analysis suite (documented and freely available for download online http://surfer.nmr.mgh.harvard.edu/). This segmentation of the pre-implant MRI with SPM12 provides us with both the tissue probability maps (i.e. gray, white, and cerebrospinal fluid (CSF) probabilities) and the indexed-binary representations (i.e., either gray, white, CSF, bone, or soft tissues). This information allowed us to reject electrodes not located in the brain. Third, the post-implant CT scan was coregistered to the pre-implant MRI via a rigid affine transformation and the pre-implant MRI was registered to the MNI template (MNI 152 Linear), via a linear and a non-linear transformation from SPM12 methods (Penny et al., 2011), through the FieldTrip toolbox (Oostenveld et al., 2011). Fourth, applying the corresponding transformations, we mapped channel locations to the pre-implant MRI brain that was labeled using the volume-based Human Brainnetome Atlas (Fan et al., 2016).

### Signal preprocessing

Continous signal was filtered using 1) a notch filter at 50Hz and harmonics up to 300Hz to remove power line artifacts, 2) a bandpass filter between 0.5Hz and 300Hz.

To define artifacted channel we used the broadband (raw) signal delimited on the experimental task recording. Channels with a variance greater than 2*IQR (interquartile range, i.e. a non-parametric estimate of the standard deviation) were tagged as artifacted channels (6% of the channels). Channels defined as artifacted were excluded from subsequent analysis.

The continuous signal during the task was then epoched from -0.2 to 14s relative to the onset of the first stimulus (repetition) of each trial. Generating 24 or 48 epochs depending on the number of sessions each patient had completed. A baseline correction was applied from -0.1 to 0s. The first 500ms were discarded from the epoched data to avoid the activity burst generated at the onset of the stimulus.

The continuous signal from the resting state session was also epoched in seventeen 14-second non-overlapping epochs.

### Power spectral density

The power spectral density (PSD) computation was conducted for each channels, with the MNE-python function Epochs.compute_psd, trial-by-trial (epochs) in a range of 125 frequencies, logarithmically scaled, ranging from 0.5 to 125 Hz. Six canonical frequency bands were investigated by averaging the power spectra between their respective boundaries: delta (1-4 Hz), theta (4-8 Hz), alpha (8-13 Hz), beta (13-30 Hz), low gamma (30-50 Hz) and high frequency activity or HFa (70-125 Hz).

### Global effect

The global effect of the task (versus period of rest) was computed on each frequency band by subtracting first the mean resting state activity from the mean experimental activity and then dividing by the mean resting state activity. This approach has the advantage of centering the magnitude before expressing it in percentage (Mercier et al., 2022). For each frequency band and channel, the statistical difference between task activity and the baseline (resting) was estimated with permutation tests (N=1000) using the SciPy library.

### Coupling behavioural and neurophysiological data

Behavioural speech data and neurophysiological data were jointly analysed using three approaches. In the first approach a two step procedure was used. First, for each frequency band, channel and trial, we computed the mean power. Then, for each frequency band and channel, we computed the correlation across trials using a non-parametric correlation metric (Spearman) between the power and the verbal coordination index (VCI between VP and patient speech). A rho value was thus attributed to each channel and frequency band. The significance was assessed using a permutation approach, similar to the one used for the global effect (see above). In the second approach, we computed the phase-amplitude coupling (PAC) between speech phase and HFa. More precisely, we used two measures for the phase: 1) the phase of the speech envelope of the VP, corresponding to the speech input, 2) the instantaneous phase difference between VP and patient phases, corresponding to the instantaneous coordination of participant and virtual partner (see Figure 2B). As for power, we used the high frequency activity from 70 to 125 Hz (considered as a proxy for activity population-level spiking activity of neurons ; Buzsáki et al., 2012). The computation was performed using Tensorpac, an open-source Python toolbox for tensor-based phase-amplitude coupling (PAC) measurement in electrophysiological brain signals (Combrisson et al. 2020). In the third approach, we assessed whether the phase-amplitude relationship (or coupling) depends upon the anticipatory (negative delays) or compensatory (positive delays) behaviour between the VO and the patients’ speech. We computed the average delay in each trial using a cross-correlation approach on speech signals (between patient and VP) with the MATLAB function *xcorr*. A median split (patient-specific ; average median split = 0ms, average sd = 24ms) was applied to conserve a sufficient amount of data, classifying trials below the median as « anticipatory behaviour » and trials above the median as « compensatory behaviour ». Then we conducted the phase-amplitude coupling analyses on positive and negative trials separately.

### Clustering analysis

A spatial unsupervised clustering analysis (k-means) was conducted on all significant channels, separately for the global effect (task versus rest) and brain-behaviour correlation analyses. Precisely, we used the silhouette score method on the k-means result (Rousseeuw, 1987; Shahapure & Nicholas, 2020). This provides a measure of consistency within clusters of data (or alternatively the goodness of clusters separation). Scores were computed for difference numbers of clusters (from two to ten). The highest silhouette score indicates the optimal number of clusters. Clustering and slihouette scores were computed using the Scikit-learn’s Kmeans and silhouette score function (Pedregosa et al., 2011). The statistical difference between spatial clustering in global effect and brain-behaviour correlation was estimated with linear model using the R function *lm* (stat package), post-hoc comparisons were corrected for multiple comparisons using the Tukey test (lsmeans R package ; Lenth, 2016). The statistical difference between clustering in global effect and behaviour correlation across the number of clusters was estimated using permutation tests (N=1000) by computing the silhouette score difference between the two conditions.

## Results

### Speech coordinative behaviour

The present synchronous speech task allowed us to create a more or less predictable context of coordination, with the objective of obtaining a wide range of coordination variability. Indeed, the intrinsic nature of the task (speech coordination) on one side and the variable coupling parameter of the VP on the other, require continuous subtle adjustments of the participants speech production. The degree of coordination between speech signals (VP and participant) was assessed at the syllabic level computing the phase locking value between the speech temporal envelopes, a measure of the strength of the interaction (or coupling) of two signals. This metric, the verbal coordination index (VCI), is a proxy of the quality of the performance in coordinating speech production with the VP. Overall, the coordinative behaviour was affected, for both controls and patients, by the coupling parameters of the model with a better coordination when the VP was set to synchronize with participants compared to when it was set to speak with a 180° syllabic shift (controls: t ratio = 4.55, p = <.0001; patients: t ratio = 6.53, p = <.0001, see boxplots in Figure 2C). This produced, as desired, a rather large coupling variability (controls: range : 0.06 - 0.84 ; mean : 0.49 ; median : 0.49 ; patients range : 0.11 - 0.94 ; mean : 0.55 ; median : 0.54, see Fig 2C). This variability was also present at the individual level (see, for patients only, Figure S1). Nonetheless, while variable, the coordinative behaviour was significantly better than chance for every patient (see Figure S1).

### Synchronous speech strongly activates the language network from delta to high gamma range

To investigate the sensitivity of synchronous speech in generating spectrally-resolved neural responses, we first analyzed the neural responses in both a spatially and spectrally resolved manner with respect to a resting-state baseline condition. In the left hemisphere, neural responses are present in all six canonical frequency bands, from the delta range (1-4 Hz) up to high frequency activity (HFa, 70-125 Hz, see Fig 3A) with medium to large modulation (increase or decrease) in activity compared to baseline (Figure 3A). More precisely, while theta, alpha and beta bands show massive desynchronisation, in particular in the STG BA41/42 (primary auditory cortex), STG BA22 (secondary auditory cortex), and IFG BA44 (Broca’s area), the low gamma and HFa bands are dominated by power increase in particular in the auditory cortex (STG BA41/42) and in the inferior frontal gyrus (IFG BA44). This modulation between synchronization and desynchronization across frequencies was significant (F(5)=6.42, p<.001 ; estimated with linear model using the R function *lm*).

**Figure 3.**
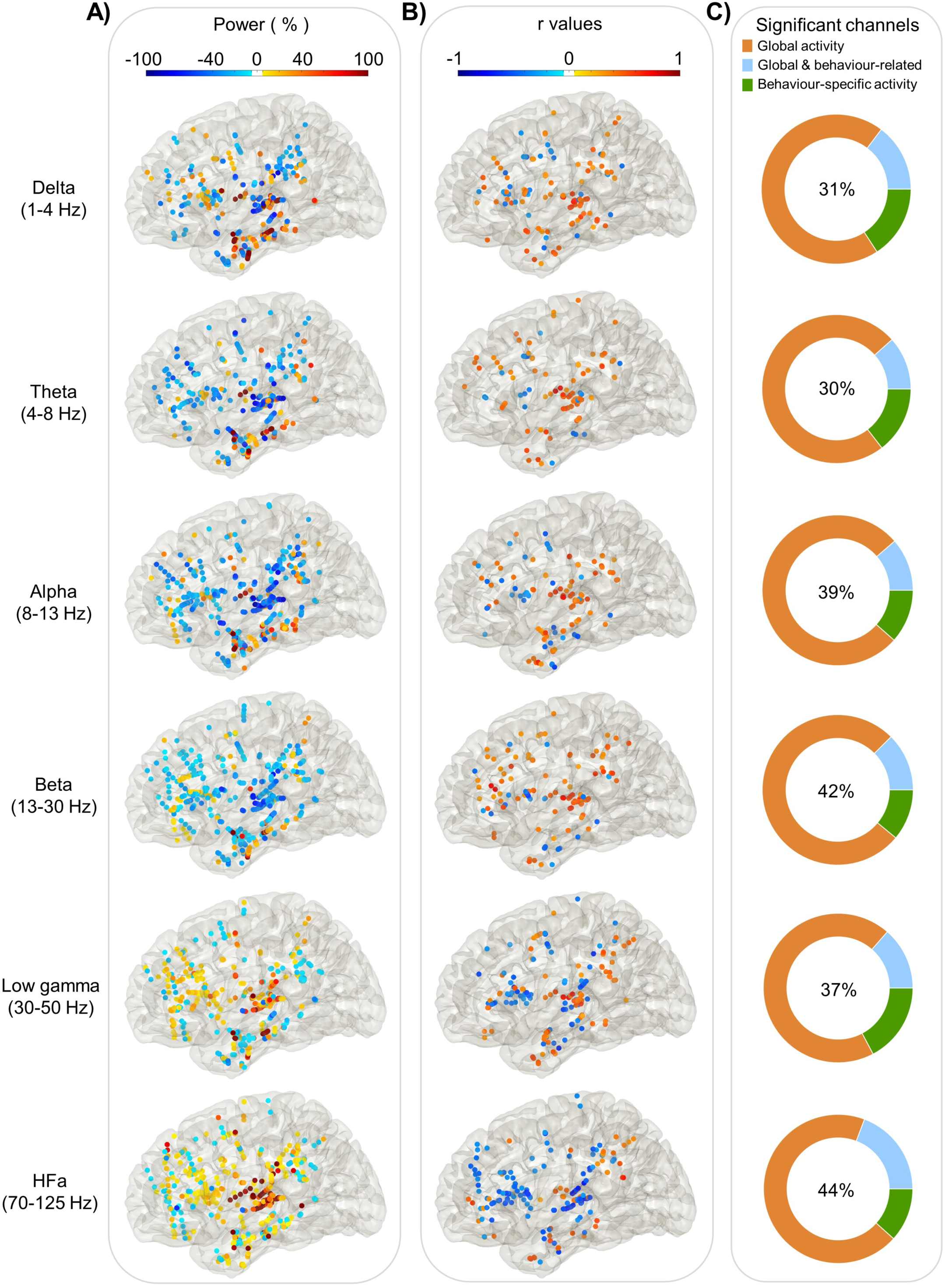
Power spectrum analyses and correlation with verbal coordination index (left hemisphere). Each dot represents a channel where a significant effect was found either on **(A)** Global activity (Task versus Rest) for each frequency band. The activity is expressed in % of power change compared to resting; or on **(B)** Behaviour-related activity: r values of the Spearman correlation across trials between the iEEG power and the verbal coordination index (VCI). **(C)** The proportion of channels where a significant effect was found: in the task vs rest (orange), in the brain-behaviour correlation (green) or for both comparison (blue). The percentage in the center indicates the overall proportion of significant channels from the three categories with respect to the total number of channels.

As expected, the whole left hemisphere language network is strongly involved, including both dorsal and ventral pathways (Fig 3A). More precisely, in the temporal lobe the superior, middle and inferior temporal gyri, in the parietal lobe the inferior parietal lobule (IPL) and in the frontal lobe the inferior frontal gyrus (IFG) and the middle frontal gyrus (MFG). Similar results are observed in the right hemisphere, neural responses are present across all six frequency bands with medium to large modulation in activity compared to baseline (Figure S2A) in the same regions. Desynchronizations are present in the theta, alpha and beta bands while the low gamma and HFa bands show power increases.

### The brain needs behaviour: global vs behaviour-specific neural activity

Comparing the overall activity during the task to activity during rest gives a broad view of the network involved in verbal coordination. However, as stated above, the task was conceived to engender a rather wide range of verbal coordination across trials for each participant. This variety of coordinative behaviours allows to explore the link between verbal and simultaneous neural activity, by computing the correlation across trials between the verbal coordination index and the mean power (see methods). This analysis allows us to estimate the extent to which neural activity in each frequency band is modulated as a function of the quality of verbal coordination.

Figure 3B shows the significant r values (Spearman correlation) for each frequency band in the left hemisphere (see Figure S2B for the right hemisphere). The first observation is a gradual transition in the direction of correlations as we move up frequency bands, from positive correlations at low frequencies to negative ones at high frequencies (F(5)=2.68, p=.02). This effect, present in both hemispheres, mimics the reversed desynchronization/synchronization process in low and high frequency bands reported above. In other words, while in the low frequency bands stronger desynchronization goes along with weaker verbal coordination, in the HFa stronger activity is associated with weaker verbal coordination.

Importantly, compared to the global activity (task vs rest, Fig 3A), the neural spatial profile of the behaviour-related activity (Fig 3B) is more clustered, in the left hemisphere. Indeed, silhouette scores are systematically higher for behaviour-related activity compared to global activity, indicating greater clustering consistency across frequency bands (t(106)=7.79, p<.001, see Figure S3). Moreover, silhouette scores are maximal, in particular for HFa, for five clusters (p < .001), located in the IFG BA44, the IPL BA40 and the STG BA41/42 and BA22 (see Figure S3).

Comparing in both hemispheres global activity and behaviour-related activities shows that, for each frequency band, approximately 2/3 of the channels are only significant in the global activity (Figure 3C, orange part). Of the remaining third, half of the significant channels show a modulation of power compared to baseline that also significantly correlates with the quality of behavioural synchronisation (Figure 3C, blue part). The other half are only visible in the brain-behaviour correlation analysis (Figure 3C, green part, behaviour-specific).

### Spectral profiles in the language network are nuanced by behaviour

In order to further explore the brain-behaviour relation in an anatomically language-relevant network, we focused on high frequency activity (HFa) and on those regions (ROI) of the dorsal pathway recorded in at least 7 patients for the left hemisphere and at least 5 patients for the right hemisphere : STG BA41/42 (primary auditory cortex), STG BA22 (secondary auditory cortex), IPL BA40 (inferior parietal gyrus) and IFG BA44 (inferior frontal gyrus, Broca’s region). To balance data completeness and statistical power, we included only brain regions recorded in at least 7 patients (∼44% of the cohort) for the left hemisphere and at least 5 patients for the right hemisphere (∼31% of the cohort), ensuring sufficient representation while minimizing biases due to sparse data. Within each ROI, using spearman correlation, we quantified the link between neural activity and the degree of behavioural coordination. Figure 4B (left hemisphere) shows a dramatic decrease of HFa along the dorsal pathway. While left-STG BA41/42 presents the strongest power increase (compared to baseline), it shows no significant correlation with verbal coordination (t(28)=-1.81, p=.08 ; Student’s T test, FDR correction). By contrast, the left-STG BA22 shows both a significant power increase in the HFa and a significant negative correlation between HFa and behaviour (i.e., VCI) (t(29)=-4.40, p<.0001 ; Student’s T test, FDR correction), marking a fine distinction between primary and secondary auditory cortex. Finally, the brain-behaviour correlation is maximal in the left-IFG BA44 (t(26)=-5.60, p<.0001 ; Student’s T test, FDR correction).

**Figure 4.**
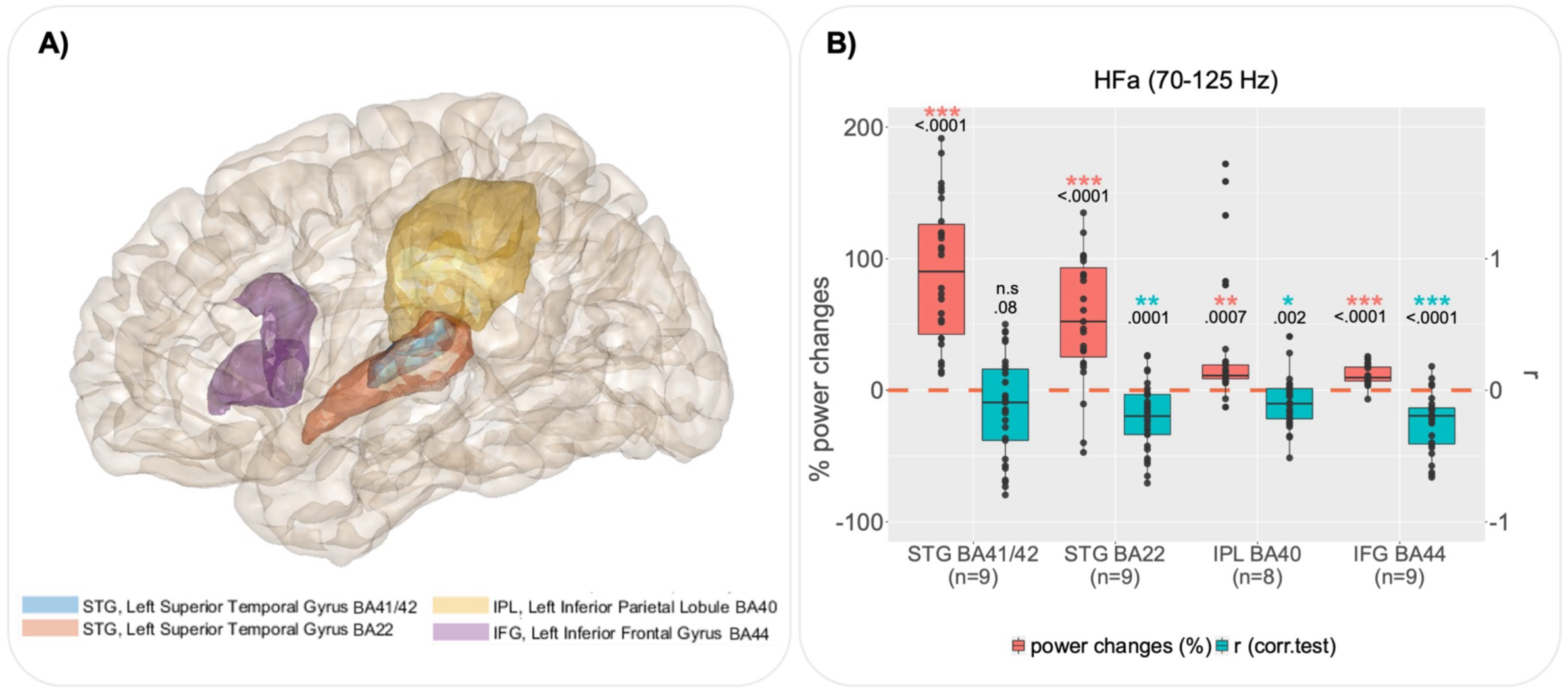
Group analysis by regions of interest for the left hemisphere. **(A)** Regions of interest (ROI) defined according to the cluster analysis (see Figure S3), the delimitation of regions is based on the Brainnetome atlas. **(B)** For each ROI, boxplots illustrate, in red, channels with significant global power changes (HFa, task vs rest) and, in blue, their corresponding r values (correlation between HFa power and verbal coordination index, VCI). Red and blue stars indicate a significant difference from a null distribution. Dots represent independent iEEG channels. The « n » below each region of interest specifies the number of patients. STG : superior temporal gyrus ; IPL : inferior parietal lobule ; IFG : inferior frontal gyrus ; BA : brodmann area.

The decrease in HFa along the dorsal pathway is replicated in the right hemisphere (Figure S4). However, while both the right STG BA41/42 and STG BA22 present a power increase (compared to baseline) — with a stronger increase for the STG BA41/42 — neither shows a significant correlation with verbal coordination (t(45)=-1.65, p=.1 ; t(8)=-0.67, p=.5 ; Student’s T test, FDR correction). By contrast, results in the right IFG BA44 are similar to the one observed in the left hemisphere with a significant power increase associated with a negative brain-behaviour correlation (t(17)=-3.11, p=.01 ; Student’s T test, FDR correction).

### The IFG is sensitive to speech coordination dynamics

To model the temporal dynamics of the relation between verbal coordination and neural activity we conducted a behaviour-brain phase amplitude coupling (PAC) analysis at the single trial level. That is, we used the power of high frequency neural activity (HFa: 70-125 Hz) and the low-frequency phase of the behaviour; the latter being either the phase of the speech signals of the virtual partner (VP) or the verbal coordination dynamics (i.e., the phase difference between VP and speaker).

When looking at the analysis of the left hemisphere (Figure 5A), coupling is strongest as expected in the auditory regions (STG BA41/42 and STG BA22), but it is also present in the left IPL and IFG. Notably, when comparing – within the regions of interest previously described – the PAC with the virtual partner speech and the PAC with the phase difference, the coupling relationship changes when moving along the dorsal pathway: a stronger coupling in the auditory regions with the speech input, no difference between speech and coordination dynamics in the IPL and a stronger coupling for the coordinative dynamics compared to speech signal in the IFG (Figure 5B). When looking at the right hemisphere, we observe the same changes in the coupling relationship when moving along the dorsal pathway, except that no difference between speech and coordination dynamics is present in the right secondary auditory regions (STG BA22 ; Figure S5). Similar results were obtained when using the phase of the patient speech rather than the VP speech (as a control analysis). Finally, in order to assess whether the phase-amplitude relationship is different for anticipatory (negative delays) and compensatory (positive delays) behaviour between the VO and the patients’ speech we assessed the difference between PAC in trials with negative and positive delays (Figure S6). Although there seems to be a trend in the left IFG with anticipatory behaviour (negative lags) being associated to stronger neural coupling, this difference did not reach significance.

**Figure 5.**
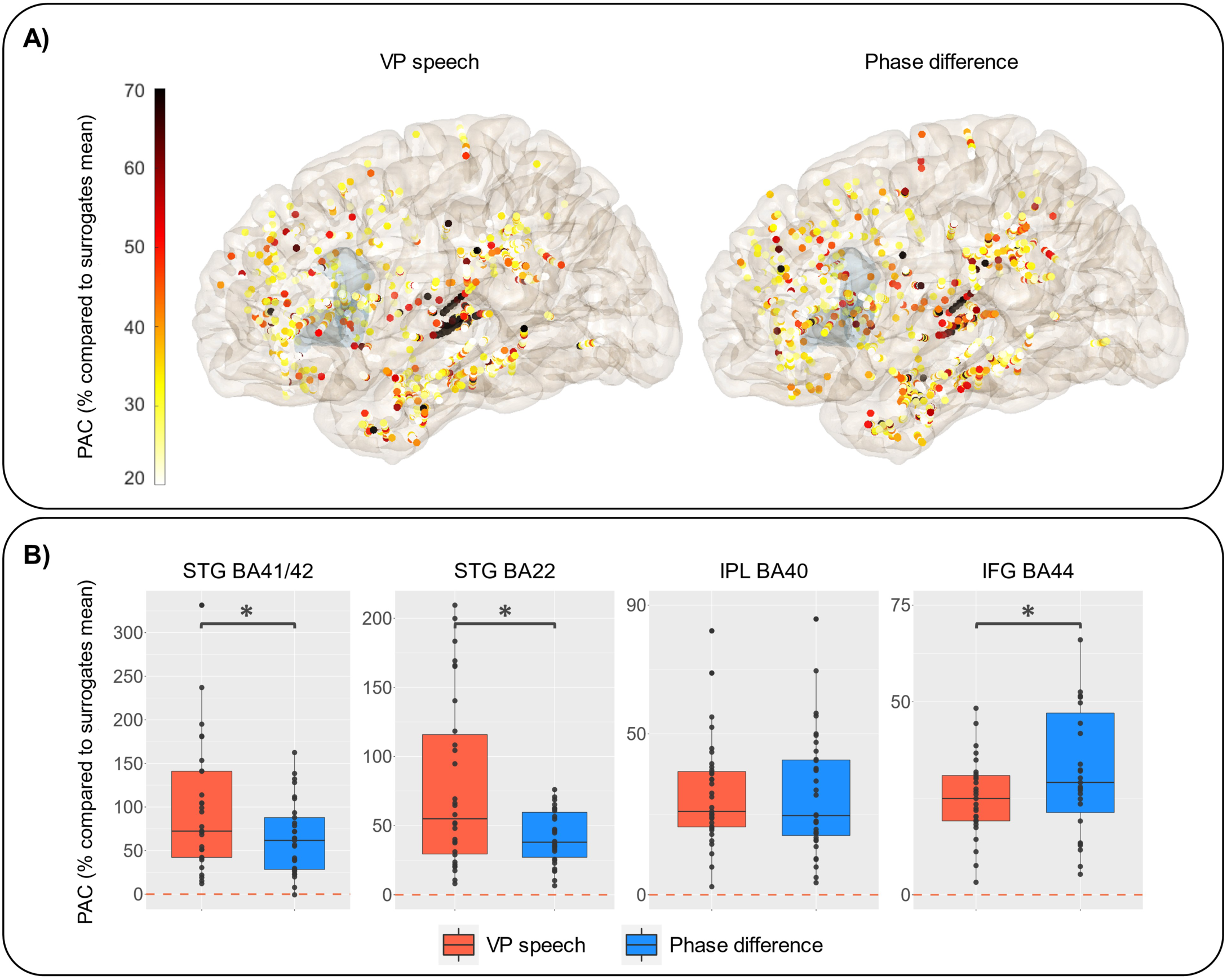
Phase-amplitude coupling between virtual partner speech signal or coordination dynamics and high frequency activity (HFa). **A)** Representation of the increase in PAC expressed in % compared to surrogates mean when using the virtual partner speech (left) or the coordination dynamics (phase difference between VP and patient, right). The shaded (slight blue) area corresponds to the location of the IFG BA44**. B)** PAC values for VP (in red) and phase difference (in blue) by regions of interest. Statistical difference between the two types of PAC is calculated using paired Wilcoxon’s test (STG BA41/42 : p=.01 ; STG BA22 : p=.004 ; IPL BA40 : p=.6 ; IFG BA44 : p=.02). Y-axis range has been adjusted to better illustrate the contrast between VP speech and coordination dynamics.

## Discussion

In this study, we investigated speech coordinative adjustments using a novel interactive synchronous speech task (Lancia et al., 2017). To assess the relation between speech coordination and neural dynamics, we capitalized on the excellent spatiotemporal sensitivity of human stereotactic recordings (sEEG) from 16 patients with drug-resistant epilepsy while they produced short sentences along with a virtual partner (VP). Critically, the virtual partner was able to coordinate and adapt its verbal production in real-time with those of the participants, thus enabling to create a variable context of coordination yielding a broad distribution of phase delays between participant and VP productions. Several interesting findings can be emphasized. Firstly, the task involving both speech production and perception is efficient to highlight both the dorsal and ventral pathways, from low to high frequency activity (Figure 3). Secondly, spectral profiles of neural responses in the language network are nuanced when combined with behavioural data, highlighting the fact that some regions are involved in the task in a general manner, while others are sensitive to the quality of verbal coordination (Figure 3). Thirdly, high-frequency activity in left secondary auditory regions shows a stronger sensitivity to behaviour (coordination success) compared to left primary auditory regions (Figure 4). Finally, the high-frequency activity of the IFG BA44 (bilaterally) specifically indexes the online coordinative adjustments that are continuously required to compensate for deviations from synchronisation (Figure 5).

### Left secondary auditory regions are more sensitive to coordinative behaviour

In the left hemisphere, the increase of high frequency activity (HFa as compared to rest) in both the STG BA41/42 (primary auditory cortex) and STG BA22 (secondary auditory cortex) is associated with different cognitive functions. Indeed, when considering whether HFa correlates with the behavioural coordination index – here the phase locking value between the patient and speaker speech – only STG BA22 shows a significant (inverse) correlation with a reduced HFa in strongly coordinated trials. This spatial distinction made between primary and secondary auditory regions in terms of their sensitivity to task demands has already been observed in the auditory cortex, particularly in the HFa (Nourski, 2017). The use of auditory target detection paradigms – phonemic categorisation (Chang et al., 2011), tone detection (Steinschneider et al., 2014), and semantic categorisation tasks (Nourski et al., 2015) – where participants are asked to press a button when they hear the target, revealed task-dependent modulations only in the STG (posterolateral superior temporal gyrus or PLST) and not in Heschl’s gyrus. In these studies, modulations corresponded to increases in activity in response to targets compared to non-targets, which have been interpreted in terms of selective attention or a bias toward behaviourally relevant stimuli (Petkov et al., 2004 ; Mesgarani & Chang, 2012). Here we extend these findings to a more complex and interactive context.

The observed negative correlation between verbal coordination and high-frequency activity (HFa) in left STG BA22 suggests a suppression of neural responses as the degree of behavioural synchrony increases. This result is reminiscent of findings on speaker-induced suppression (SIS), where neural activity in auditory cortex decreases during self-generated speech compared to externally-generated speech (Meekings & Scott, 2021; Niziolek et al., 2013). However, our paradigm differs from traditional SIS studies in two critical ways: (1) the speaker’s own voice is always present and predictable from the forward model, and (2) no passive listening condition was included. Therefore, our findings cannot be directly equated with the original SIS effect. Instead, we propose that the suppression observed here reflects a SIS-related phenomenon specific to the synchronous speech context. Synchronous speech requires simultaneous monitoring of self- and externally-generated speech, a task that is both attentionally demanding and coordinative. This aligns with evidence from Ozker et al. (2022, 2024), showing that the same neural populations in STG exhibit SIS and heightened responses to feedback perturbations. These findings suggest that SIS and speech monitoring are related processes, where suppressing responses to self-generated speech facilitates error detection. In our study, suppression of HFa as coordination increases may reflect reduced prediction errors due to closer alignment between perceived and produced speech signals. Conversely, increased HFa during poor coordination may signify greater mismatch, consistent with prediction error theories (Houde & Nagarajan, 2011; Friston et al., 2020). Furthermore, when self- and externally-generated speech signals are temporally and phonetically congruent, participants may perceive external speech as their own. This echoes the “rubber voice” effect, where external speech resembling self-produced feedback is perceived as self-generated (Zheng et al., 2011; Lind et al., 2014; Franken et al., 2021). While this interpretation remains speculative, future studies could incorporate subjective reports to investigate this phenomenon in more detail.

Furthermore, the absence of correlation in the right STG BA22 (Figure S4) seems in first stance to challenge influential speech production models (e.g. Guenther & Hickok, 2016) that propose that the right hemisphere is involved in feedback control. However, one needs to consider the the task at stake heavily relied upon temporal mismatches and adjustments. In this context, the left-lateralized sensitivity to verbal coordination reminds of the works of Floegel and colleagues (2020, 2023) suggesting that both hemispheres are involved depending on the type of error: the right auditory association cortex monitoring preferentially spectral speech features and the left auditory association cortex monitoring preferentially temporal speech features. Nonetheless, the right temporal lobe seems to be sensitive to speech coordinative behaviour, confirming previous findings using fMRI (Jasmin et al., 2016) and thus showing that the right hemisphere has an important role to play in this type of tasks (e.g. Jasmin et al., 2016).

### Inferior frontal gyrus (BA44) as a site of speech coordination/planning in dynamic context

Our results highlight the involvement of the inferior frontal gyrus (IFG) bilaterally, in particular the BA44 region, in speech coordination. First, trials with a weak verbal coordination (VCI) are accompanied by more prominent high frequency activity (HFa, Fig.4; Fig.S4). Second, when considering the within-trial time-resolved dynamics, the phase-amplitude coupling (PAC) reveals a tight relation between the low frequency behavioural dynamics (phase) and the modulation of high-frequency neural activity (amplitude, Fig.5B ; Fig.S5). This relation is strongest when considering the phase adjustments rather than the phase of speech of the VP per se : larger deviations in verbal coordination are accompanied by increase in HFa. Additionally, we also tested for potential effects of different asynchronies (i.e., temporal delay) between the participant’s speech and that of the virtual partner but found no significant differences (Fig.S6). While lack of delay-effect does not permit to conclude about the sensitivity of BA44 to absolute timing of the partner’s speech, its neural dynamics are linked to the ongoing process of resolving phase deviations and maintaining synchrony.

These findings are in line with the importance of higher-level frontal mechanisms for behavioural flexibility and their role in the hierarchical generative models underlying speech perception and production (Cope et al., 2017). More precisely, they are in line with works redefining the role of Broca’s area (BA44 and BA45) in speech production associating it more to speech planning rather than articulation per se (Flinker et al., 2015; Basilakos et al., 2018). Indeed, electrodes covering Broca’s area show a greatest activity before the onset of articulation and not during speech production. This has been interpreted in favour of a role at a prearticulatory stage rather than an « on-line » coordination of the speech articulators at least in picture naming (Schuhmann et al., 2009), word repetition (Flinker et al., 2015; Ferpozzi et al., 2018) and turn-taking (Castellucci et al., 2022). According to these studies Broca’s area may be a « functional gate » at a preacticulatory stage, allowing the phonetic translation before speech articulation (Ferpozzi et al., 2018). Our use of a synchronous speech task allows to refine this view by showing that these prearticulatory commands are of continuous rather than discrete nature. In other terms, the discrete (on-off) and ignition-like behaviour of neuronal populations in Broca’s area gating prearticulatory commands before speech may be due to the discrete nature of the tasks used to assess speech production. Notably, picture naming, word repetition, word reading and even turn-onsets imply that speech production is preceded by a silent period during which the speaker listens to speech or watches pictures. By contrast, the synchronous speech task requires continuous temporal adjustments of verbal productions in order to reach synchronisation with the virtual partner. Relatedly, the involvement of IFG in accurate speech timing has been previously shown via thermal manipulation (Long et al., 2016).

Of note, temporal adjustments (prediction error corrections) are also needed for fluent speech in general, beyond synchronous speech, and give rise to the rhythmic nature of speech. Temporal adjustments possibly take advantage of the auditory input and are referred to as audio-motor interactions that can be modeled as a coupled oscillator (Poeppel & Assaneo, 2020). Interestingly, during speech perception, the coupling of theta and gamma bands in the auditory cortex reflects tracking of slow speech fluctuations to spiking gamma (Morillon et al., 2010; Morillon et al., 2012; Hyafil et al., 2015; Lizarazu et al. 2019; Oganian & Chang, 2019; Leonard et al., 2024), similar to what we describe in the auditory cortex. By contrast, in the inferior frontal gyrus, the coupling in the high-frequency activity is strongest with the input-output phase difference (input of the VP - output of the speaker), a metric that could possibly reflect the amount of error in the internal computation to reach optimal coordination. This indicates that this region could have an implication in the optimisation of the predictive and coordinative behaviour required by the task. This well fits with the anatomical connectivity that has been described between Broca’s and Wernicke’s territory via the long segment of the arcuate fasciculus, possibly setting the base for a mapping from speech representations in Wernicke’s area to the predictive proprioceptive adjustments processed in Broca’s area (Catani & Ffytche, 2005; Oestreich et al., 2018). This also aligns with broader theories on the relationship between perception and action, such as predictive coding and active inference, which propose shared sensory prediction mechanism and neural computational architecture for both processes (Friston et al., 2017, 2020).

Finally, while the case of synchronous speech may seem quite far away from real-life conversational contexts, the models describing language interaction consider that listeners covertly imitate the speaker’s speech and timely construct a representation of the underlying communicative intention which allows early fluent turn-taking (Pickering & Gambi, 2018; Levinson, 2016). Moreover, synchronous speech has recently gained interest in the neuroscience field due to important results showing a relation between anatomo-functional features and synchronization abilities. More precisely, Assaneo and collaborators (2019) used a spontaneous speech synchronization (SSS) test wherein participants produce the syllable /tah/ while listening to a random syllable sequence of predefined pace. The authors identified two groups of participants (high and low synchronisers) characterised by their ability to naturally synchronise their productions more or less easily with the auditory stimulus. Importantly, the ability to synchronise correlates to the degree of lateralization of the arcuate fasciculus, that connects the inferior frontal gyrus and the auditory temporal regions, high synchronizers showing greater lateralization to the left than the low synchronizers. The more this structural connectivity of the arcuate fasciculus is lateralised in the left hemisphere, the more the activity of the IFG is synchronised with the envelope of the audio stimulus of the SSS-test, during a passive listening task. A major limitation of EEG and MEG studies is that they are very sensitive to speech production artefacts, which is not the case of iEEG. Thus, the full dynamics of speech interaction are difficult to assess with surface recordings. Our findings extend these results in several manners. First, speech production is not limited to a single syllable, but to complex utterances. Secondly, the input auditory stimulus is not preset but adapts and changes behaviour in real-time as a function of the dynamic of the « dyad », here patient-virtual partner. Thirdly, and most importantly, iEEG recording allowing speech artefact-free data, we could extend the relation between coordination abilities and anatomical circuitry of the IFG to the neural dynamics of this same region, showing that it plays an important role in the temporal adjustment of speech that are necessary to coordinate to external speech.

To conclude, the present study illustrates the possibility and interest of using a fully dynamic, adaptive and interactive language task to gather deeper understanding of the subtending neural dynamics involved in speech perception, production as well as their interaction. It is worth noting that the influence of specific speech units, such as consonants versus vowels, on speech coordination remains to be explored. In non-interactive contexts, participants show greater sensitivity during the production of stressed vowels, possibly reflecting heightened attentional or motor adjustments (Oschkinat & Hoole, 2022; Li & Lancia, 2024). In this study, the VP’s adaptation relies on sensitivity to spectral cues, particularly phonetic transitions, with some (e.g., formant transitions) being more salient than others. However, how these effects manifest in an interactive setting remains an open question, as both interlocutors continuously adjust their speech in real time. Future studies could investigate whether coordination signals, such as phase resets, preferentially align with specific parts of the syllable.

## Supplementary Material

**Figure S1.**
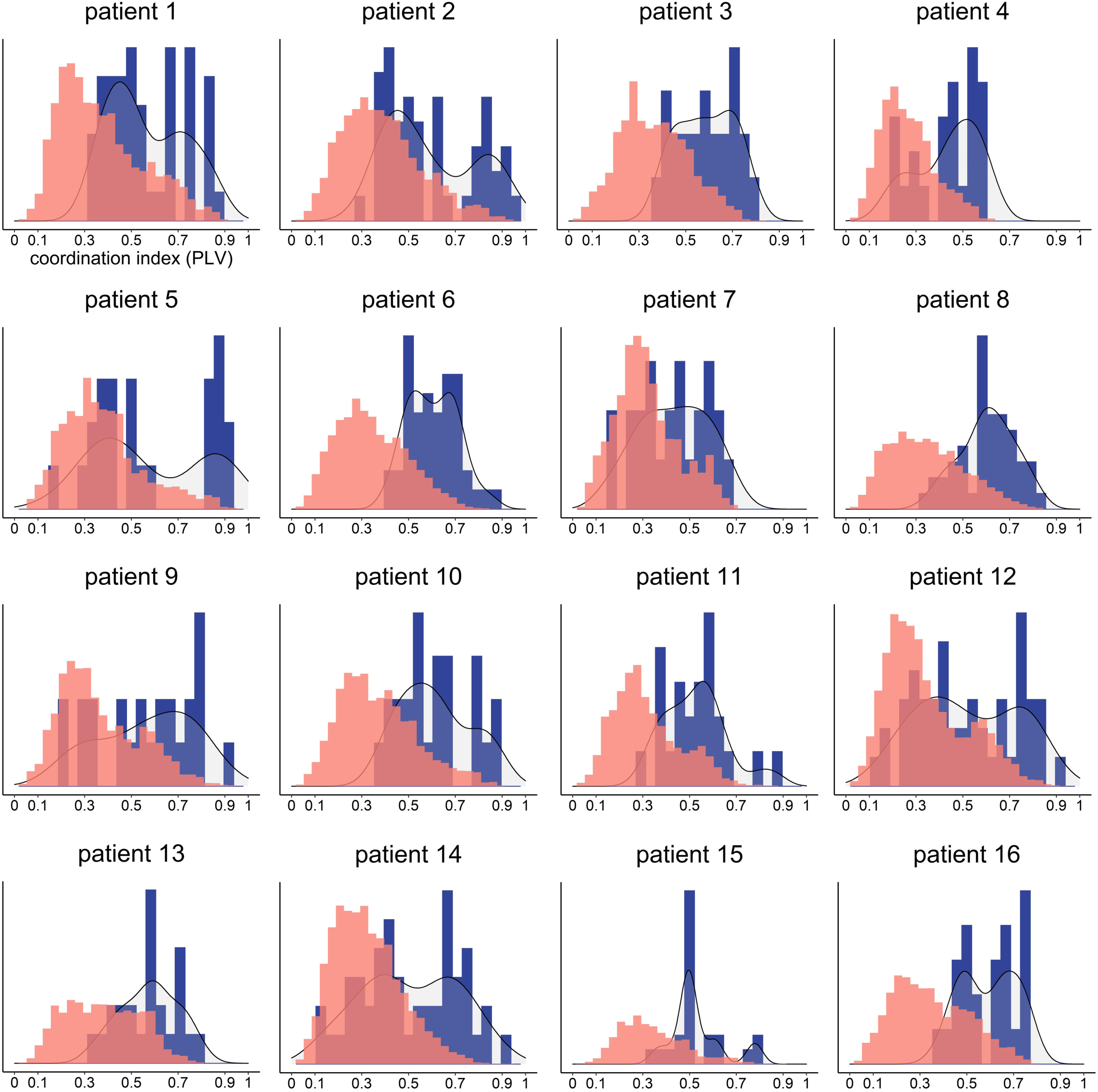
Distribution of verbal coordination index for each patient. (PLV between patient’s speech and VP speech). For each of the sixteen patients, this figure depicts the histogram of the coordination index for all trials (in blue) as well as the null distribution (random phase shift) computed using 500 permutations per trial (in red).

**Figure S2.**
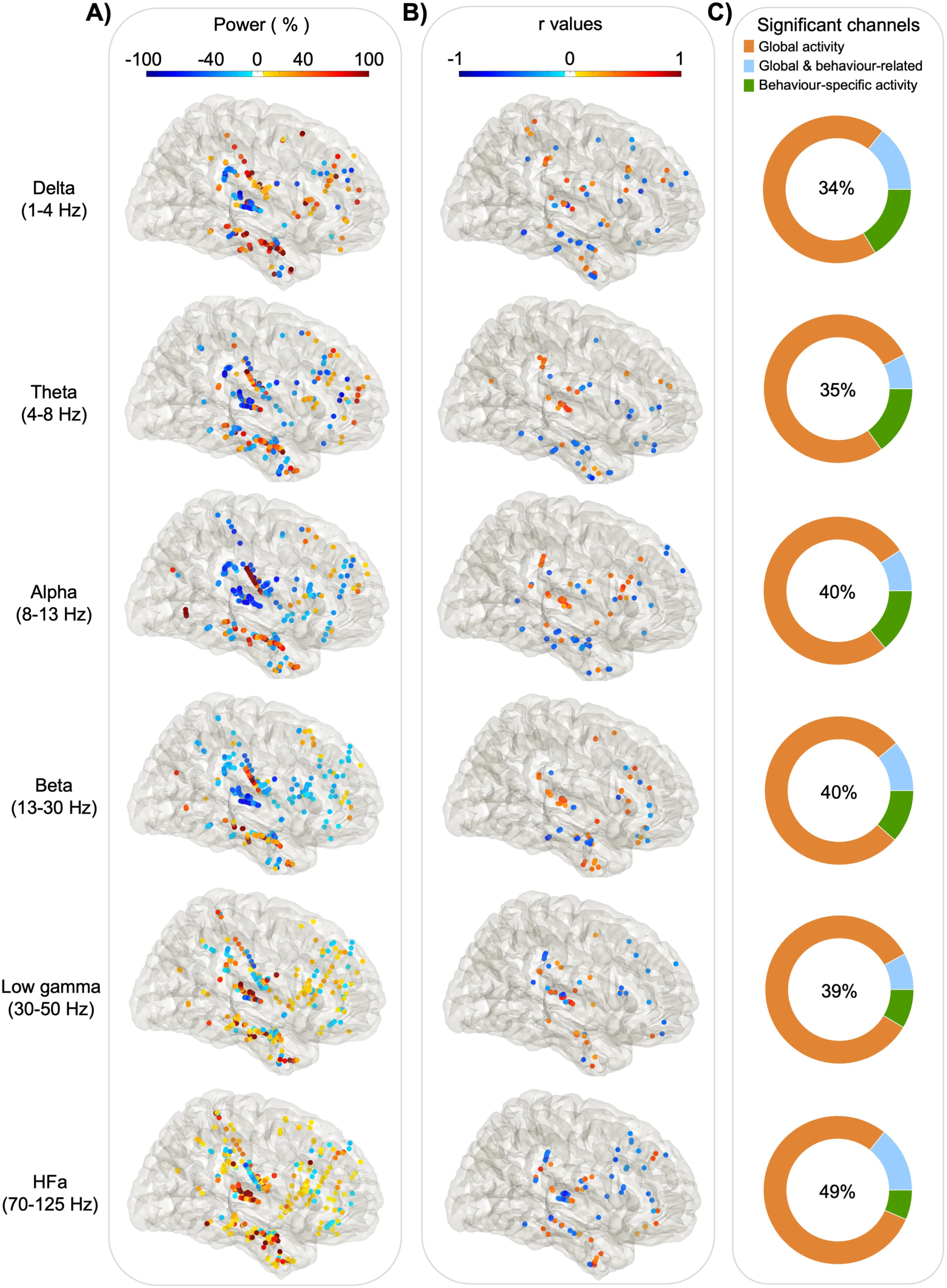
Power spectrum analyses and correlation with verbal coordination index (right hemisphere). Each dot represents a channel where a significant effect was found either on **(A)** Global activity (Task versus Rest) for each frequency band. The activity is expressed in % of power change compared to resting; or on **(B)** Behaviour-related activity: r values of the Spearman correlation across trials between the iEEG power and the verbal coordination index (VCI). **(C)** The proportion of channels where a significant effect was found: in the task vs rest (orange), in the brain-behaviour correlation (green) or for both comparison (blue). The percentage in the center indicates the overall proportion of significant channels from the three categories with respect to the total number of channels.

**Figure S3.**
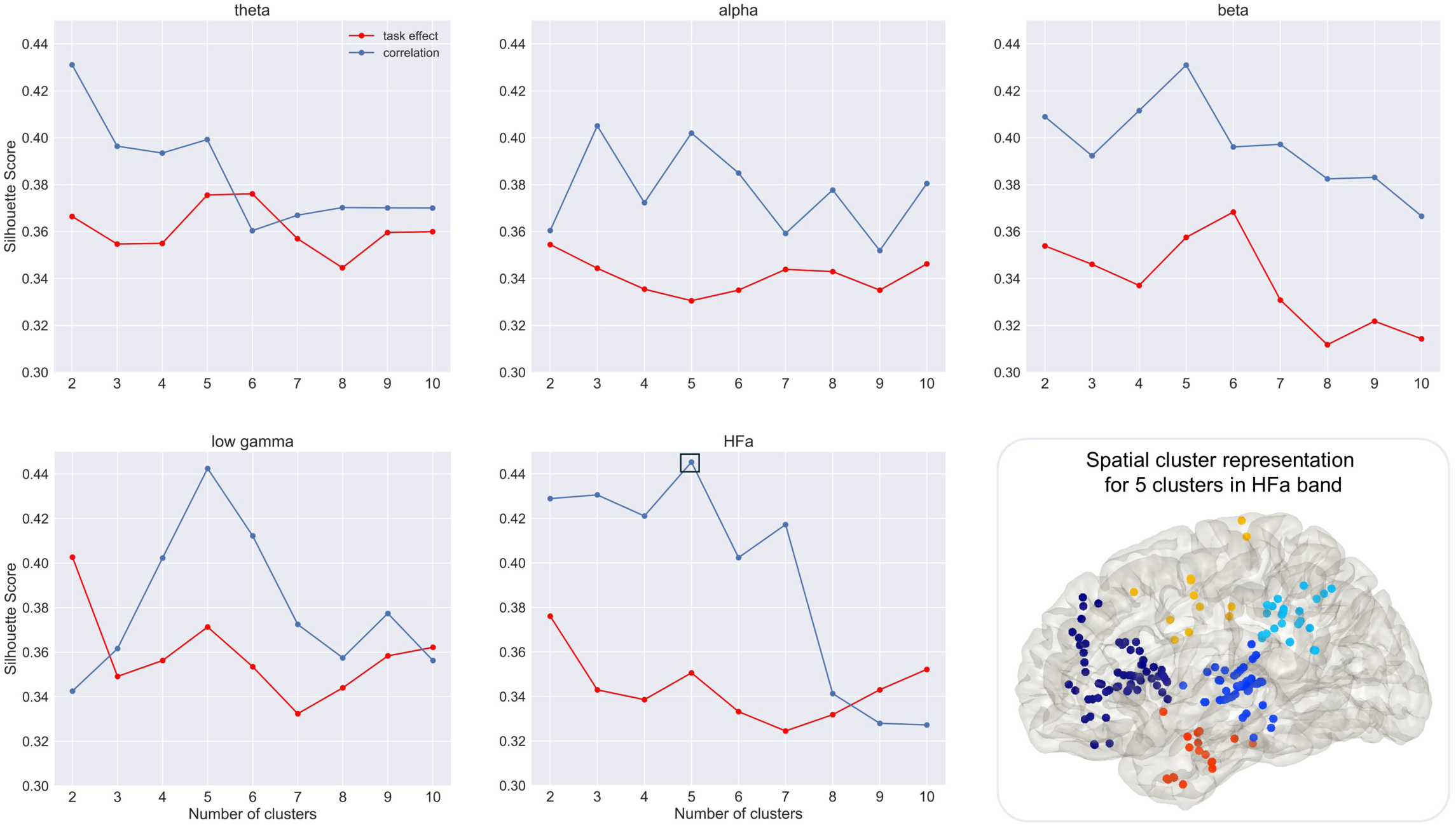
Cluster analysis (silhouette score). Illustration of the mean silhouette scores according to the number of clusters for global activity (in red) and behaviour-related activity (correlation between power changes and coordination index, in blue). The highest silhouette score was obtained for five clusters in the HFa range for behaviour-related activity (framed in full black square). Bottom right: spatial cluster representation for the highest mean silhouette score value in HFa range.

**Figure S4.**
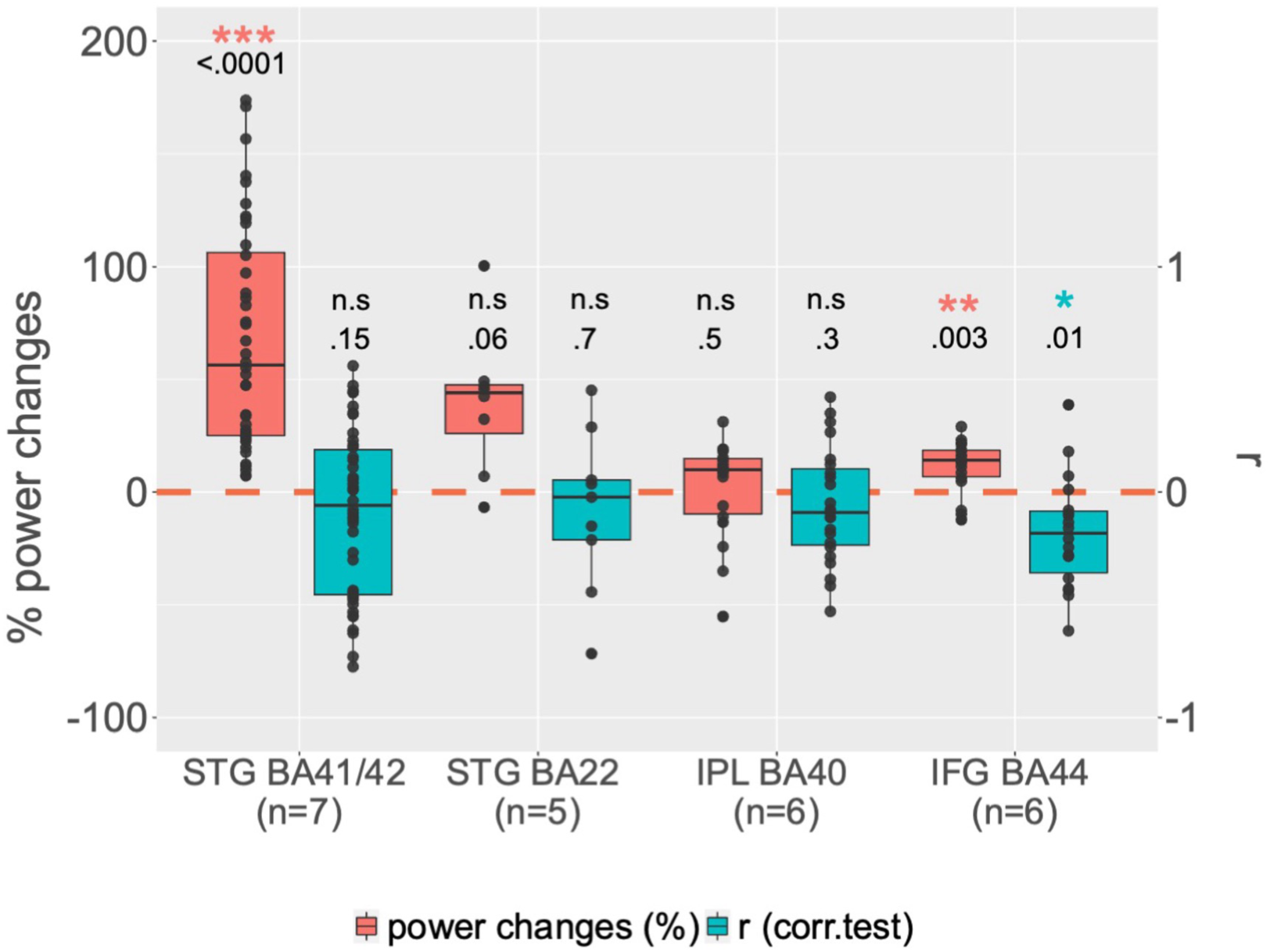
Group analysis by regions of interest (right hemisphere). For each ROI, boxplots illustrate, in red, channels with significant global power changes (HFa, task vs rest) and, in blue, their corresponding r values (correlation between HFa power and verbal coordination index, VCI). Red and blue stars indicate a significant difference from a null distribution. Dots represent independent iEEG channels. The « n » below each region of interest specifies the number of patients. STG : superior temporal gyrus ; IPL : inferior parietal lobule ; IFG : inferior frontal gyrus ; BA : Brodmann area.

**Figure S5.**
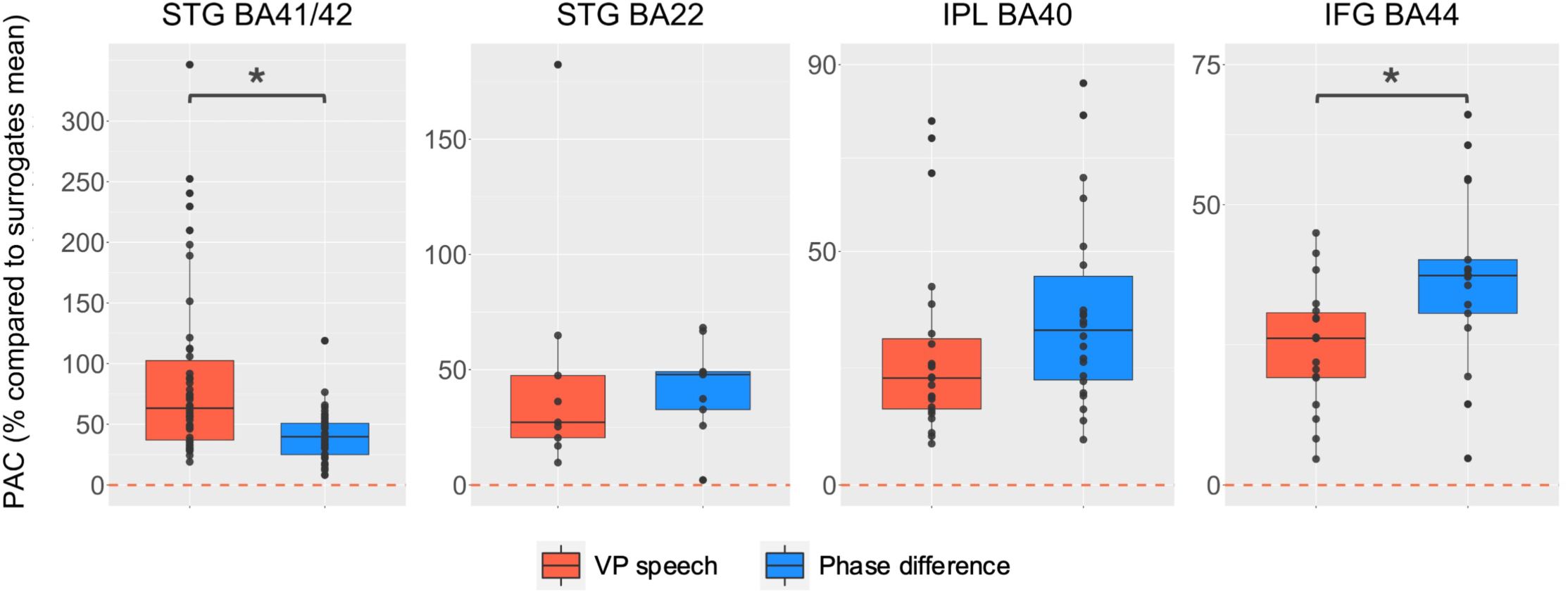
Phase-Amplitude Coupling analyses by region of interest (right hemisphere). PAC expressed in % compared to surrogates when using as phase the virtual partner speech (in red) or the coordination dynamic (phase difference between VP and participant, in blue) and as amplitude the high frequency activity. Statistical difference between the two different types of PAC is calculated using paired wilcoxon test (STG BA41/42 : p<0.0001 ; STG BA22 : p=0.9 ; IPL BA40 : p=0.2 ; IFG BA44 : p=0.01). Y-axis range has been adjusted to better illustrate the contrast between VP speech and coordination dynamic.

**Figure S6.**
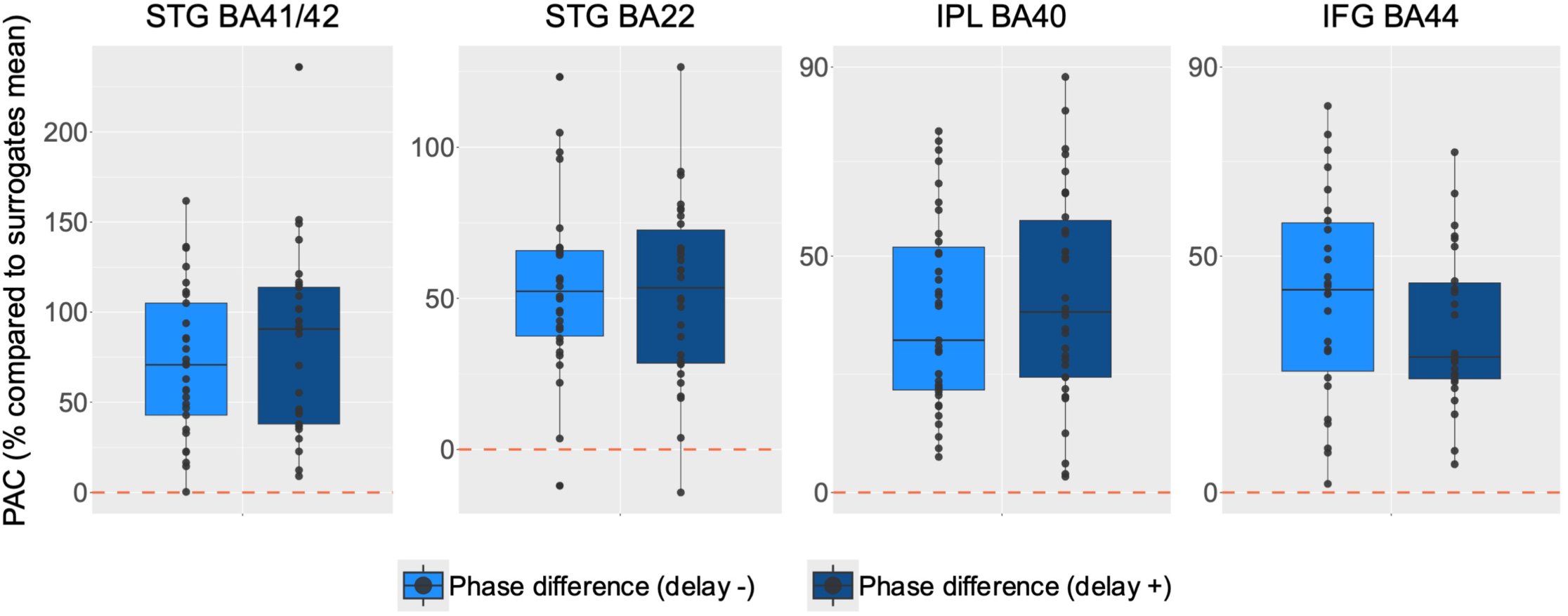
Phase-Amplitude Coupling according to the behavioural delay (left hemisphere). PAC expressed in % compared to surrogates when using as phase the coordination dynamic (phase difference between VP and participant) and as amplitude the high frequency activity. Comparison between PAC on trials with negative and positive delays (see “Coupling behavioural and neurophysiological data” section in Materials and methods). Please note that the y-axis range has been adjusted per panel.

## Acknowledgments

We thank all patients for their willingful participation and all the personnel from the epileptology unit.

## Author contributions

I.S.M, L.L and D.S. designed research; L.L. wrote the code for the experimental setup ; I.S.M., L.L, A.T. and M.M. acquired data; I.S.M, M.M., L.L and D.S. analyzed data; I.S.M and D.S. wrote the paper; I.S.M, A.T., M.M., B.M., L.L and D.S. edited the paper.

## Data availability statement

The conditions of our ethics approval do not permit public archiving of anonymised study data. Readers seeking access to the data should contact Dr. Daniele Schön. Access will be granted to named individuals in accordance with ethical procedures governing the reuse of clinical data, including completion of a formal data sharing agreement.

## Code availability statement

Data analyses were performed using custom scripts in Matlab & Python, and will be available upon publication on Github.

